# discoal: flexible coalescent simulations with selection

**DOI:** 10.1101/063453

**Authors:** Andrew D. Kern, Daniel R. Schrider

**Affiliations:** Department of Genetics, Rutgers University, Piscataway, NJ, 08854, USA; Human Genetics Institute of New Jersey, Rutgers University, Piscataway, NJ, 08554, USA

## Abstract

**Summary:** Here we describe `discoal`, a coalescent simulator able to generate population samples that include selective sweeps in a feature-rich, flexible manner. `discoal` can perform simulations conditioning on the fixation of an allele due to drift or either hard or soft selective sweeps—even those occurring a large genetic distance away from the simulated locus. `discoal` can simulate sweeps with recurrent mutation to the adaptive allele, recombination, and gene conversion, under nonequilibrium demographic histories and without specifying an allele frequency trajectory in advance.

**Availability and Implementation:** Availability and implementation: discoal is implemented in the C programming language. Source code is freely available on GitHub (https://github.com/kernlab/discoal_multipop) under a GNU General Public License.

**Supplementary information:** Supplementary Figures and Text are appended below

## 1 Introduction

The coalescent process, which describes the stochastic genealogy of a sample of alleles from a natural population, has become the dominant paradigm for both theory and empirical analysis in population genetics over the past three decades (Hudson, 1990; Nordborg, 2001). Indeed the great efficiency of coalescent simulations, relative to forward-in-time population genetic simulations, allows researchers to perform statistical inference of population genetic parameters directly from simulations (e.g. Pritchard *et al.*, 1999; Wall *et al.*, 2002). Such efforts have mostly focused on inferring demographic parameters such as population size histories or migration rates (e.g. Excoffier *et al.*, 2013; Fagundes *et al.*, 2007; Naduvilezhath *et al.*, 2011), and accordingly the majority of coalescent simulation software has focused on being flexible with respect to the range of demographic histories they can model (Hudson, 2002).

Increasingly, population genomics as a field has recognized the role that natural selection has played in shaping patterns of variation in natural populations (e.g. Langley *et al.*, 2012; Sella *et al.*, 2009). The coalescent again provides a convenient framework for modeling the action of selection on population genetic samples. In particular good coalescent approximations to both selective sweeps (Braverman *et al.*, 1995; Kaplan *et al.*, 1989; Kim and Stephan, 2002) and balancing selection (Kaplan *et al.*, 1988) have been described analytically. While that is so there are relatively few software packages that flexibly model the coalescent with selection; notable exceptions include msms (Ewing and Hermisson, 2010) and mbs (Teshima and Innan, 2009). Here we introduce `discoal`, a coalescent simulator that incorporates genic selection in a flexible manner. `discoal` is able to simulate standard demographic events, such as population splits and mergers, instantaneous population size changes, admixture, and migration, along with selective sweeps that occur either at the locus sampled or in neighboring loci. `discoal` is further capable of simulating both hard and soft selective sweeps (Pennings and Hermisson, 2006), while conditioning on the fixation of the selected allele. In addition `discoal` simulates coalescent samples that are conditional on the fixation of a neutral mutation.

## 2 Methods

In settings without selection, `discoal` simulates realizations of the Ancestral Recombination Graph (ARG) in a manner similar to Hudson’s algorithm, where recombination break points are discrete and the number of sites possible for recombination is a parameter of the simulation. In the context of a selective sweep, `discoal` uses the structured coales-cent approach of Braverman et al. (1995), where during a “sweep phase” rates of coalescence depend on the current frequency of the selected allele as it spreads (forward in time) through the population. Conditioning on selected allele trajectories is performed in `discoal` one of two possible ways as chosen by the user: sweeps can follow deterministic trajectories or instead can be stochastic through the use of a conditional diffusion approach both using a model of genic (i.e. haploid) selection (Coop and Griffiths, 2004; Kim mid Stephan, 2002). Neutral fixation trajectories can also be simulated by `discoal`. To accomplish this we use a stochastic jump process between small time steps, *δt*, to simulate the allele frequency, *p*, moving back in time from frequency 1.0 to frequency 1/2*N* such that the frequency of the neutral mutation in the next step, *p′*, is given by

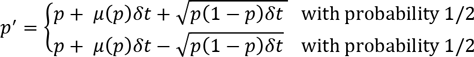

with *¼*(*p*) = −*p* (Ewens, 2004). This trajectory routine was tested for accuracy by calculating the expected time to fixation and by comparing simulation results to those from Tajima (1990). To model stochastic hard sweeps the selected allele frequency, *p*, is again modeled as a jump process as above, however in this case *¼*(*p*) is given by

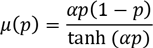

where *α* = 2*Ns* is the population scaled selection coefficient. We verified the accuracy of our simulations conditioning on the recent fixation of hard sweeps by comparing them to simulations from msms, finding very close agreement (Supplementary Fig. S1). To simulate soft sweeps we introduce an additional parameter to the model, *f_0_*, the frequency at which the allele came under directional selection. To generate trajectories from this model, we simulate a stochastic selection trajectory back in time until the frequency *f_0_*, is reached, and then switch over to a neutral fixation trajectory until absorption as in Przeworski *et al.* (2005). `discoal` can also simulate soft sweeps arising from recurrent mutation to the adaptive allele at a rate specified by the user.

Simulation of conditional trajectories via jump processes allows us to model selection in the context of instantaneous population size changes naturally, as the coalescent is a Markovian process: if a population size boundary interrupts a sweep phase of the coalescent, event probabilities (i.e. of coalescence, recombination, etc.) are simply reset to reflect the new population size and the simulation continues. In this manner `discoal` can simulate coalescent genealogies that condition on selective sweeps or neutral fixations in the presence population size changes. This functionality is absent from msms, and mbs can perform such simulations only if the user supplies an appropriate set of frequency trajectories ahead of time, `discoal` can also approximate genealogies of populations experiencing continuous population size changes by simulating a series of discrete changes. We verified the accuracy of our jump process trajectories with changing population size through comparison to forward time, single site simulations and comparisons to summary statistics generated using SLiM (Messer, 2013; see Supplementary Figs. S2-S5).

`discoal` allows for an arbitrary number of subpopulations, but limits selective sweeps to only occur in a single population. Further we do not allow migration between other populations and the population undergoing a sweep during the sweep phase of the simulation. We make this assumption in order to avoid modeling sweep phase dynamics in multiple populations concurrently, `discoal` is also uniquely capable of simulating coalescent genealogies with sweeps that occur within the sampled region as well as those occurring some specified recombination distance away (c.f. Braverman *et al.*, 1995).

As an example of the kinds of analyses that `discoal` enables, we have generated simulation data under each of our three conditional fixation types (hard sweep, soft sweep, and neutral) and examined patterns of linked population genetic variation at increasing recombination distances from the site of the fixation. Supplementary Fig. S6 shows the decay towards equilibrium values for five population genetic summary statistics, `discoal` is available for download on GitHub (https://github.com/kem-lab/discoal_multipop). The manual for `discoal` is available on the GitHub repository and Supplementary Text S1.

## Funding

A.D.K. was supported in part by NIH award no. R01GM078204. D.R.S. was partially supported by the National Institutes of Health under Ruth L. Kirsch-stein National Research Service Award F32 GM105231.

*Conflict of Interest:* none declared.

